# Emergent Traveling Waves in Neural Circuits

**DOI:** 10.64898/2026.01.08.698281

**Authors:** Navid Shervani-Tabar, Scott L. Brincat, Mikael Lundqvist, Earl K. Miller

## Abstract

Artificial neural networks achieve striking performance but do not capture a prominent neural property. Cortical activity is organized by structured spatiotemporal dynamics, traveling waves (TW), which have been implicated in a wide range of functions. Existing computational models often rely on hand-crafted connectivity and imposed dynamics, offering insight into their impact but less into how the waves naturally emerge in biological circuits. Here, we found that TWs emerge in models under biologically plausible constraints (spatially organized, directionally biased connectivity). Under an empirical neural manifold constraint, these wiring principles naturally emerge to support traveling wave dynamics in the recurrent model. We further show that wave propagation provides a robust mechanism for maintaining working memory in the presence of visual distractors. We compared these model predictions to non-human primate prefrontal cortex recordings, revealing a similar mechanism. Together, these results advance our understanding of traveling waves as a substrate for cognition and offer a framework for mechanistic accounts of cortical computation.

## Introduction

Traveling waves are synchronous, spatially organized patterns of neural activation that propagate across laminar neural tissues [1, 2, 3]. These waves have emerged as a fundamental feature of large-scale brain dynamics, arising spontaneously [3, 4, 5] or in response to external stimuli [6, 7, 5]. They are observed across sensory [3, 4, 6, 7, 5], motor [8, 9], and cognitive areas [10, 11]. As they traverse neural circuits, traveling waves transiently modulate neuronal excitability [12, 13], coordinate activity across distributed networks [14, 15], and support perception [3, 16, 17, 18] and memory [10, 19, 11]. Notably, a wave’s spatial configuration, temporal dynamics, and direction of propagation can encode functional and behavioral significance [19, 3, 15, 11, 13, 14, 20, 21]. Collectively, these findings underscore traveling waves as a dynamic substrate for spatiotemporal computation in neural circuits [14].

Integrating traveling waves into recurrent neural networks (RNNs) [22] offers a biologically grounded path to spatiotemporal computation in neural circuit models. Benigno et al. [23] create waves in a reservoir computing model by placing a complex-valued recurrent layer on a topographic grid with local Gaussian connectivity. Each connection has a distance-dependent delay that mimics axonal conduction and promotes coherent wave propagation. However, using fixed, isotropic distance functions for recurrent connections and delays prevents the network from learning task-specific recurrent structure. This limits the generalization and interpretability of neural dynamics. Similarly, Keller & Welling [24] and Jacobs et al. [25] place a coupled-oscillatory RNN [26] on a spatial lattice and use shared local convolutional kernels in place of dense recurrence. Nearest-neighbor, one-step recurrence enforces local wave propagation, but restricts the development of task-specific, long-range, or location-dependent recurrent structures. Likewise, Keller et al. [27] impose a wave equation directly on the RNN’s update rule by using the discrete shift operator instead of dense recurrence. This operator supports only uniform transport within each channel, limiting spatial heterogeneity in wave dynamics and making it difficult to express patterns such as rotational waves.

Here, we introduce a data-constrained framework where traveling waves emerge in spatially embedded recurrent networks without prescribing wave equations or rigid lattice topology. The approach pairs an anatomical prior, i.e., units arranged on a two-dimensional sheet with distance-dependent coupling, with a functional prior that aligns the network’s latent dynamics with low-dimensional manifolds recovered from non-human primate (NHP) prefrontal cortex (PFC) during working memory. Building on this, we learn the recurrent connectivity motifs associated with spatiotemporal propagation and distill them into a compact, locality-preserving parameterization. We then impose this parameterization *de novo* to construct spatially organized RNNs that, by design, can support traveling wave dynamics while remaining agnostic to any explicit wave operator or fixed grid topology. This pipeline links empirical population geometry to circuit-level design principles and provides a general recipe for assembling wave-capable recurrent models.

## Results

### Manifold-Aligned RNN exhibits emergent traveling waves

We began by modeling the neural circuit with a recurrent neural network (RNN) whose units are spatially organized. Traveling waves are spatiotemporal activity patterns that propagate across neural tissue [1, 28]. To observe these dynamics in the model, a physical embedding of the network units is necessary. In our model, units are arranged on a two-dimensional sheet that approximates the cortical surface. In the cortex, nearby neurons are more likely to be connected than distant neurons [29, 30, 31, 32, 33]. We implemented this by letting synaptic strength among units decay with inter-neuronal distance (Fig. 1a). This approach yields stronger local couplings and weaker long-range connections (see Supplementary Fig. S3). The model was trained by minimizing an objective that combines task performance error with a distance-based regularizer (see Methods; [34]).

**Figure 1:**
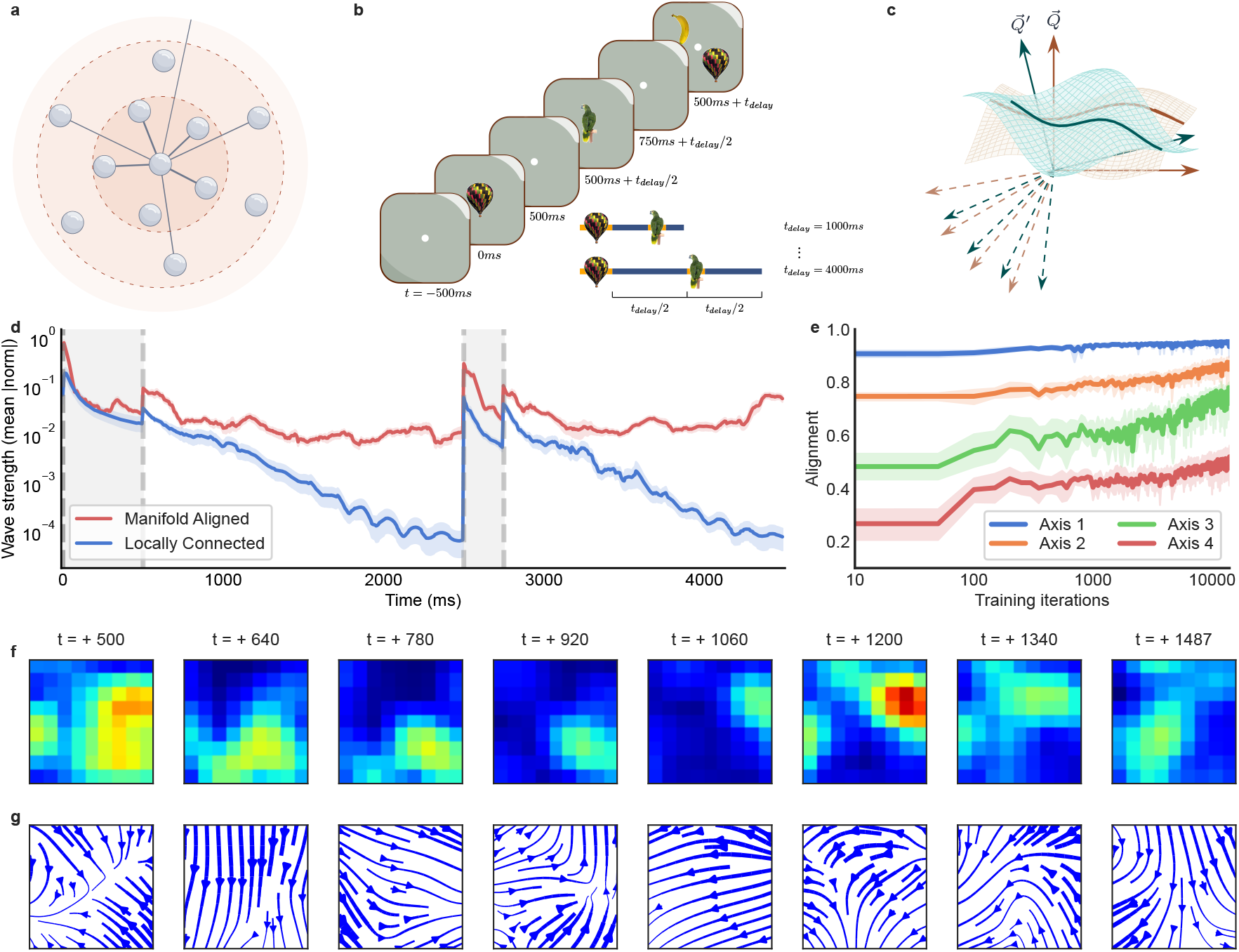
Latent manifold alignment enables persistent traveling waves in the network during working memory delay: **a**, Distance-dependent connectivity in the recurrent network. Units are arranged on a two-dimensional sheet approximating the cortical surface. Synaptic strength is more strongly regularized with increasing inter-neuronal distance (lighter shading), resulting in strong local and weak long-range couplings. Lines depict example connections from a reference unit at the center. **b**, The delayed match-to-sample (DMS) timeline used to train the network. After fixation on a central point (−500 − 0 ms; white dot), a sample image is shown for 500 ms (0 − 500 ms). A variable memory delay *t*_delay_ follows. In half of the trials, a 250-ms visual distractor appears at the midpoint of the delay. During the test, two images were displayed, and the subject/model reported the match (saccadic choice in NHP; image-coordinate readout in the model). The delay *t*_delay_ was varied across trials. The schematic depicts example timelines with variable delays (bottom right); intermediate delays are not shown for brevity. **c**, Manifold-alignment regularization. Network and empirical PFC trajectories are projected onto low-dimensional subspaces (teal and tan surfaces, respectively) with orthonormal bases ***Q***^**′**^ (model) and ***Q*** (data). Alignment is quantified by the canonical correlations between these bases. The training objective maximizes the sum of squared correlations across the axes (see Methods). **d**, Wave strength during the DMS task, quantified by the domain-averaged optical-flow magnitude, for trials with the longest delay (*t*_delay_ = 4 s). Solid lines show the mean across trials for Locally Connected (blue) and Manifold-Aligned networks (red). Shaded regions indicate plus or minus one standard deviation. Grey bands denote stimulus epochs (sample, 0 − 500 ms; mid-delay distractor, 2500 − 2750 ms). **e**, Alignment of model and empirical latent subspaces during training. Curves show canonical correlations between the model and PFC manifolds for the top four latent axes across training iterations. The shaded region denotes the 98% confidence interval (see Methods). **f-g**, Traveling wave dynamics in the Manifold-Aligned network during the memory-delay period. **f**, Snapshots from a trial with the shortest delay (*t*_delay_ = 1 s) showing wave patterns across the neuronal sheet (see Methods). **g**, Corresponding instantaneous velocity fields, estimated by optical-flow analysis. Streamlines indicate flow direction; line width encodes local speed (vector magnitude).

To investigate whether the Locally Connected RNN could exhibit task-evoked traveling waves, we trained the model on a delayed match-to-sample (DMS) task (Fig. 1b; [10]). Task-evoked traveling waves have been documented across cognitive operations, including working memory maintenance [10]. The DMS task involved presentation of a sample image, followed by a memory delay, and then by two test images. The correct response was to choose the matching image. In the middle of the memory delay, there was a visual distractor. This distractor was designed as a minor disruption to the memory.

Distance-dependent synaptic strength alone (Fig. 1a) was insufficient to sustain traveling waves. After training the RNN, we quantified wave propagation using the optical-flow algorithm [35, 36]. We applied this technique to successive activity maps from the model to infer the instantaneous velocity field on the 2-D neuronal domain (see Methods). For each trial and frame, we computed the spatial mean of the flow magnitude and averaged these traces across trials (Fig. 1d). In the Locally Connected, distance-based network, the domain-averaged flow magnitude rises at sample onset and then declines during the sample period. A smaller peak follows at the start of the delay. As the delay progresses, the magnitude decays exponentially toward zero. A visual distractor triggers another brief increase, followed by a decline once the stimulus ends. Together, these patterns indicate that distance-dependent connectivity alone is insufficient to sustain propagating waves through the memory delay.

Given this limitation, we next considered an auxiliary constraint on the population-level activity subspace. Cortical population activity is often confined to a low-dimensional subspace of the full neural space [37, 38, 39, 40]. This latent subspace, also called a neural manifold, is shaped by circuit connectivity and behavior [41, 42]. While connectivity based on distance alone provides a plausible anatomical prior (see above), the neural activity arising from this model does not necessarily span the same subspace as the population activity seen in NHP cortex [43]. We hypothesized that encouraging the model to occupy a latent subspace aligned with the empirical data from the same task would induce connectivity patterns supporting similar dynamics. To this end, we augmented training with a manifold-alignment regularizer [39, 44]. This constraint encourages the model’s latent trajectories to approximate the subspace structure observed in PFC recordings from NHP during working memory maintenance (Fig. 1c). While this alignment may not enforce the exact connectivity as the biological circuitry, it biases connectivity toward motifs that can reproduce the empirical latent dynamics.

Adding the low-dimensional manifold constraint to the training objective progressively reoriented the network’s dynamics toward that seen in the neurophysiological data (Fig. 1e). Specifically, we aligned the top four latent axes between the model and neural data. Alignment, measured by canonical correlations, improved over training: Axis 1 rose from 0.84 to near unity, Axis 4 from 0.23 to 0.61, and intermediate axes showed similar gains. During evaluation, the network received only task inputs with no extra constraints on its dynamics. In the Manifold-Aligned network, the domain-averaged flow magnitude again rose sharply at stimulus onset. Crucially, it no longer collapsed with stimulus offset and remained elevated throughout the memory delay (Fig. 1d). Trial snapshots and corresponding streamline fields show coherent wavefronts sweeping across the sheet during the delay (Fig. 1f-g).

Taken together, these results show that shaping the latent subspace structure is sufficient to sustain traveling waves during the memory delay in a spatially embedded (Locally Connected) RNN. The persistence of these waves indicates the connectivity contains structured recurrent interactions beyond simple distance-dependent decay.

### Directional, sparsely interlocked wiring supports wave propagation

We next examined how manifold alignment altered connectivity. In recurrent networks, the topology of synaptic connectivity shapes the repertoire of accessible dynamics [42, 45, 46, 47]. While structure constrains these dynamics, their expression is further gated by inputs and internal state of the network [48, 49]. To isolate the architectural features that sustained the propagation of traveling waves, we contrasted the recurrent connectivity of Manifold-Aligned vs. Locally Connected models. This comparison uncovers wiring patterns promoted by manifold alignment during training.

Both models have spatial embedding: Connections between nearby neurons are stronger than those between distant neurons. This spatial bias was confirmed by a strong negative correlation between inter-neuronal distance and connection weight in both the Manifold-Aligned and Locally Connected networks (Fig. 2a). Thus, each network was endowed with tightly knit local neighborhoods.

**Figure 2:**
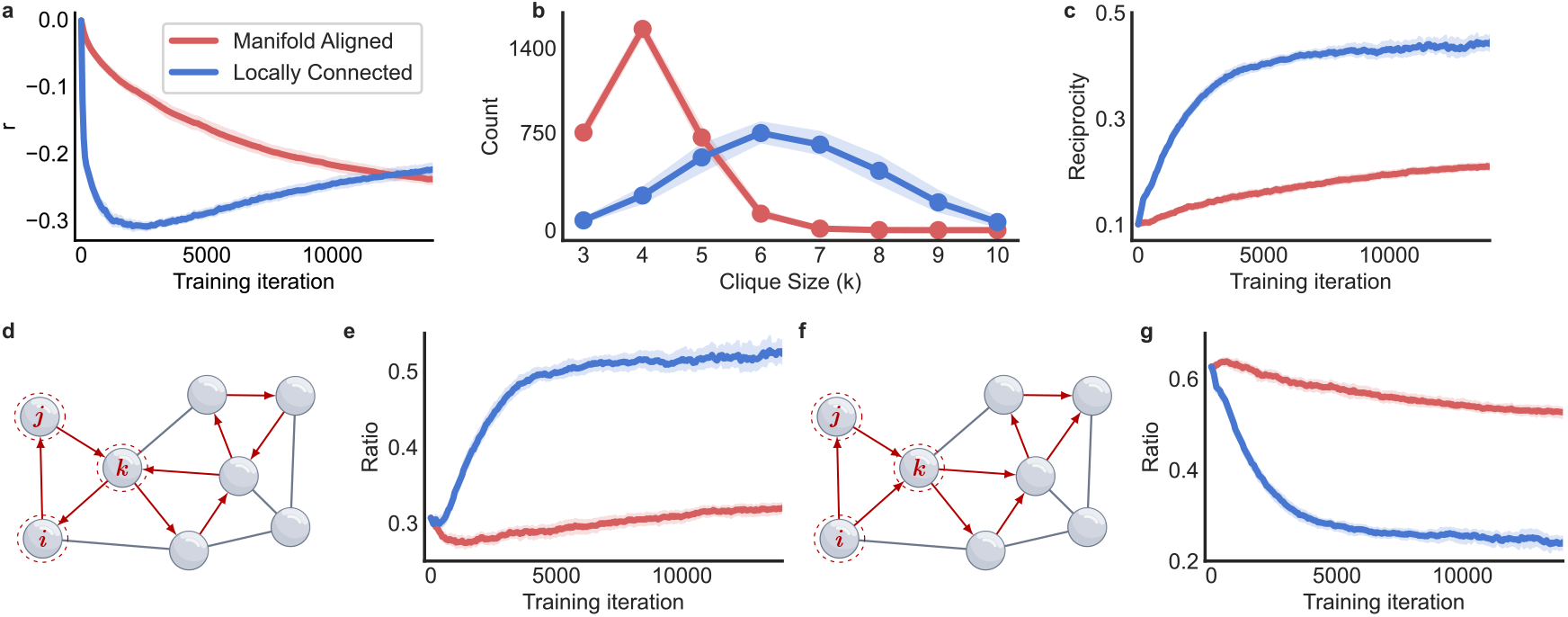
Wave propagation emerges from directionally biased networks: **a**, Correlation between inter-neuronal distance and synaptic weight across training. **b**, Distribution of maximal *k*−clique sizes in the trained network. **c**, The fraction of bidirectional edges (reciprocity) over training. **d**, Schematic of a three-node directed cycle, comprising the directed edges *i* → *j, j* → *k*, and *k* → *i*. The illustration exemplifies a network architecture enriched in such cycles. **e**, Ratio of directed cycles in the network during training. **f**, Schematic of a three-node feed-forward loop, comprising the directed edges *i* → *j, i* → *k*, and *j* → *k*. The illustration highlights a network topology with an abundance of such motifs. **g**, Relative abundance of feed-forward loop (FFL) triads, expressed as a ratio to all triads in the network over the course of training. In all plots, red denotes the Manifold-Aligned model and blue denotes the Locally Connected model. The shaded area represents the 98% confidence interval (see Methods).

However, a key distinction lies in how these neighborhoods interlock. We quantified this by measuring the population of cliques in the undirected version of the networks. This undirected view ignores the direction of synaptic connections and simply records whether any connection exists between two neurons. A maximal *k*-clique is a set of *k* all-to-all connected units (fully interconnected) that are not part of any larger clique. As clique size increases, neighborhoods become more overlapped. This affects how activations propagate in the circuit. In the Manifold-Aligned model, neighborhoods mainly form small cliques. The distribution of clique sizes peaks at *k* = 4, while larger fully interconnected neighborhoods (*k* ≥ 5) are strongly suppressed (Fig. 2b). In contrast, the Locally Connected model shows a heavy-tailed distribution of clique sizes. In this model, the structure shifts toward larger cliques, making highly interconnected neighborhoods much more prevalent.

This difference in connectivity structure has clear effects on network dynamics. In the Locally Connected network, the abundance of large cliques causes activity to spread rapidly within those clusters. This results in near-simultaneous activation of many nearby nodes. The pattern resembles a standing wave, a spatially fixed pattern of activity [1] (see Supplementary Fig. S4), rather than a traveling wave. By comparison, the Manifold-Aligned network has small cliques. This allows neighboring nodes to synchronize just enough to support a propagating wavefront. This limit on the clique size prevents immediate, widespread co-activation of the population. Thus, activity patterns in the Manifold-Aligned model manifest as a traveling wave.

The two models also differ in how the directions of synaptic connections are organized. Pairwise analysis of neurons showed that the Locally Connected model contains more bidirectional edges (connections in both directions between a pair of neurons; Fig. 2c). This indicates symmetric coupling and strong feedback. This cyclic structure is also evident in higher-order interactions: Triads (sets of three neurons) frequently form directed cycles (Fig. 2d-e), reinforcing local recurrence. By contrast, the Manifold-Aligned network is dominated by one-way connections. Triads in this model mainly form one-way chains of influence (feed-forward loop; Fig. 2f-g). In the Locally Connected network, high reciprocal coupling and cycles foster local mixing. This rapidly dissipates activity. In the Manifold-Aligned network, directed edges and feed-forward motifs relay activity sequentially. This sustains traveling waves.

Together, these comparisons show that manifold alignment shifted circuitry from recurrent, symmetric wiring to a regime that favors directed propagation. Across descriptors, from edges to motifs, the Manifold-Aligned model shows higher directionality, more feed-forward loops, fewer cycles, and lower reciprocity. These features establish a connectivity regime that sustains wave propagation in the absence of external drive.

### Spatial locality with forward-biased motifs is sufficient for sustained traveling waves

Next, we examined how the connectivity motifs identified in the preceding section are organized across the circuit to support traveling waves. Analyses of these motifs revealed that the network exhibiting traveling waves is enriched with unidirectional (one-way) rather than bidirectional (reciprocal) connections. Such directionality provides a natural substrate for sequential activation of the neurons [50, 51, 52]. However, to assemble a chain that propagates activity, successive links must cohere in sequence with aligned directions [53, 54]. Motivated by this, we asked whether the recurrent connectivity harbors a latent feedforward chain of connections.

To examine the underlying arrangement of connections, we analyzed networks based on the peak latency of neuronal activity [55]. To that end, we ordered neurons by the time of their peak firing rate and computed the mean synaptic weight as a function of rank order difference *i* − *j* across all pairs of neurons (*i, j*). In this peak-ordered view of the connectivity, the Manifold-Aligned model displayed a pronounced asymmetric peak (Fig. 3a). The asymmetry falls on the positive side of the *i* − *j* axis, meaning synapses are stronger, on average, in the forward direction. This bias indicates that earlier-firing neurons preferentially excite those that follow shortly thereafter. By contrast, the Locally Connected network showed no comparable asymmetry. Together, these observations extend the motif-based analyses by revealing a latent feed-forward scaffold embedded within the recurrent connectivity in the Manifold-Aligned model.

**Figure 3:**
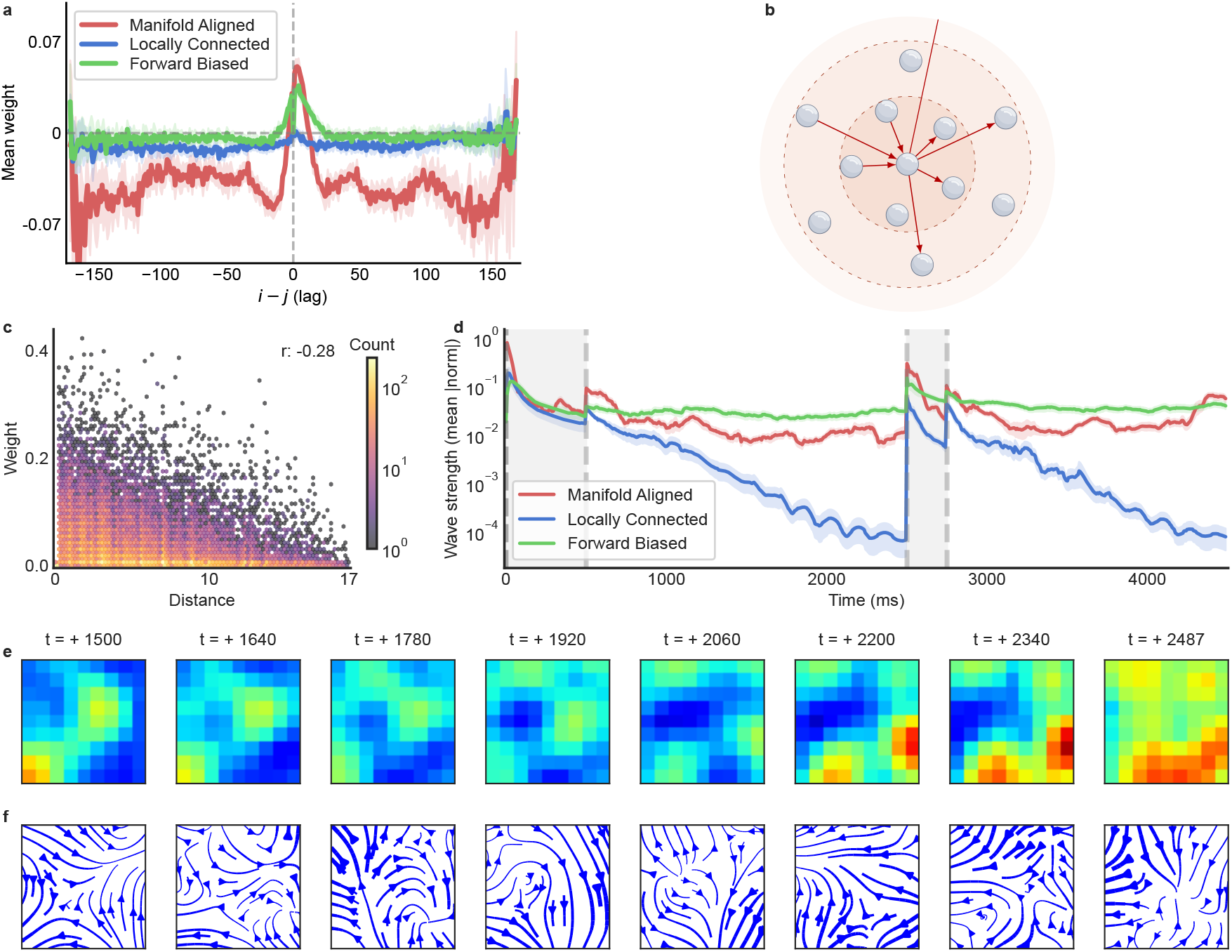
Directional bias in synaptic connectivity generates cortex-like traveling waves: **a**, Peak-ordered connectivity profile. Neurons were ranked by the time of their peak firing, and, for each rank order difference *i* − *j*, we averaged synaptic weights across all neuron pairs sharing that order difference. Positive order differences indicate “forward” connections from earlier-to later-peaking neurons. The Manifold-Aligned (red) and Forward-Biased models (green) both show pronounced, asymmetric peaks at small positive order differences, consistent with forward-directed connectivity, whereas the Locally Connected model (blue) shows little directional structure. The vertical dashed line marks zero order difference. The shaded area indicates the 98% confidence interval (see Methods). **b**, Forward-biased, distance-regularized connectivity. Units lie on a two-dimensional sheet approximating the cortex, and synaptic strength is increasingly regularized as inter-neuronal distance grows (lighter shades indicate stronger regularization). We imposed directionality by expressing the weights as a learnable similarity transform of a sub-diagonal template defined in a latent basis (see Methods). This preserves strong local and weak long-range couplings while embedding a hidden feed-forward scaffold capable of supporting traveling wave dynamics during the delay period. Arrows show example directed connections of a central reference unit. **c**, Relationship between synaptic weight magnitude and inter-neuronal distance. Each hexagon represents a bin of connections, with color indicating the number of connections per bin (log scale). The plot reveals a concentration of strong weights at short range and an extended tail of weak, long-range couplings. Across all connections, synaptic strength decreases with distance (Pearson *r* = −0.28). **d**, Temporal profile of wave strength during the memory delay in DMS task (*t*_delay_ = 4 s), quantified as the spatial mean of the optical-flow magnitude. Curves show trial-averaged trajectories for Manifold-Aligned (red), Locally Connected (blue), and Forward-Biased (green) networks; shaded areas denote one standard deviation margin across trials. Grey boxes mark stimulus periods (sample, 0 − 500 ms; mid-delay distractor, 2500 − 2750 ms). **e-f**, Propagating wave patterns in the Forward-Biased network during the memory-delay interval. **e**, Representative frames from a trial with the briefest delay (*t*_delay_ = 1 s), illustrating waves across the network (see Methods). **f**, Maps of instantaneous velocity, derived via optical flow. Streamlines depict directionality, while line thickness reflects local propagation speed.

We next asked whether introducing a latent feed-forward scaffold into an otherwise Locally Connected network is sufficient to sustain traveling waves. A naïve construction would project from each neuron predominantly to its immediate successor in index order (synapses directed from neuron *i* to neuron *i* + 1) [54, 56]. This results in a sub-diagonal weight matrix. This approach, however, proved brittle. Distance-based regularization, necessary for imposing spatial locality of the connections, clashed with this imposed order and impeded learning. To incorporate directionality while preserving spatial locality, we re-parameterized the connectivity via a learnable similarity transform. In this formulation, a sub-diagonal structure is imposed in a latent basis [57], then mapped to physical coordinates by a trainable change of basis [58]. Distance-based regularization is applied only after this transformation, on the resulting connectivity in physical coordinates. This approach thus yields a forward-biased yet spatially local connectivity pattern (Fig. 3b; see Methods). We used this architecture to test whether a latent feed-forward scaffold is sufficient to sustain traveling waves.

Using the similarity-transform parameterization, we trained the Forward-Biased network on the DMS task with distance-based regularization. In this setting, no alignment of the neural manifold was applied. When neurons were ordered by the time of their peak activity, the recurrent weight profile displayed a pronounced forward asymmetry (Fig. 3a). Synaptic strength declined with spatial distance (*r* = −0.28), consistent with the intended local connectivity (Fig. 3c). These analyses demonstrate that the parameterization preserved a latent feed-forward scaffold while enforcing local connectivity constraints.

After training, we quantified wave propagation across the spatially arranged neurons. We then computed the flow magnitude over time, averaging across both trials and the spatial domain. The Forward-Biased network showed a transient increase at sample onset that relaxed to a non-zero plateau persisting throughout the delay. A second transient was evoked by the distractor, after which the plateau recovered and remained stable (Fig. 3d). Snapshot maps and streamline visualizations corroborated this dynamic. They revealed a coherent wavefront that advanced steadily across the domain during the delay (Fig. 3e,f). By comparison, the Locally Connected network’s flow decayed toward baseline (Fig. 3d). These results establish that the Forward-Biased architecture sustains self-propagating traveling waves during memory maintenance.

### Propagation maintains working memory during distraction

We next examined how the Forward-Biased RNN dynamics shaped working memory maintenance following distraction. To perform the DMS task used here, both the model and the NHP must preserve a neural representation of the sample across the delay in face of behaviorally irrelevant distractor presented in the middle of the delay [59, 60]. We then evaluated these model predictions using *in vivo* recordings from the NHP PFC to assess their biological relevance to cortical population dynamics.

To characterize how sample and distractor information are represented during distraction, we quantified two complementary time-resolved measures of information in the population activity. First, we computed time-resolved sample-decoding accuracy, which tracked how well sample identity could be decoded from the population. Since maintaining the working memory is necessary to perform the task, the sample identity should remain decodable. We used a classifier trained to predict sample identity from neuronal firing rates. To quantify how sample information is distributed across neurons, we defined a unit-wise readout score based on decoder accuracy (see Methods). In parallel, because distractor identity is not required for task performance, we measured distractor selectivity [61] using time-resolved two-way ANOVA on firing rates, reported as 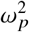 (PEV) [62]. This statistic indicated the proportion of firing rate variance attributable to the distractor after accounting for variance due to sample identity (see Methods). For each neuron, we identified the time at which it reached its peak in decoding accuracy or distractor selectivity. We then sorted neurons by these peak latencies to reveal temporal structure.

In the Forward-Biased RNN, memory-related information was propagated sequentially across the population rather than being statically maintained. Sample-decoding accuracy was not sustained within a fixed subgroup. Instead, individual neurons reached peak accuracy at different times, spanning distraction and post-distraction intervals (Fig. 4b). Sorting neurons by this peak-accuracy time produced a clear temporal cascade. This indicated the network maintained the memory by dynamically recruiting different neurons. We observed the same cascade in the NHP recordings (Fig. 4f). Distractor-evoked signals showed a similar sweep in the model (Fig. 4a) with a distinct temporal order. This ordered propagation was corroborated in PFC (Fig. 4e).

**Figure 4:**
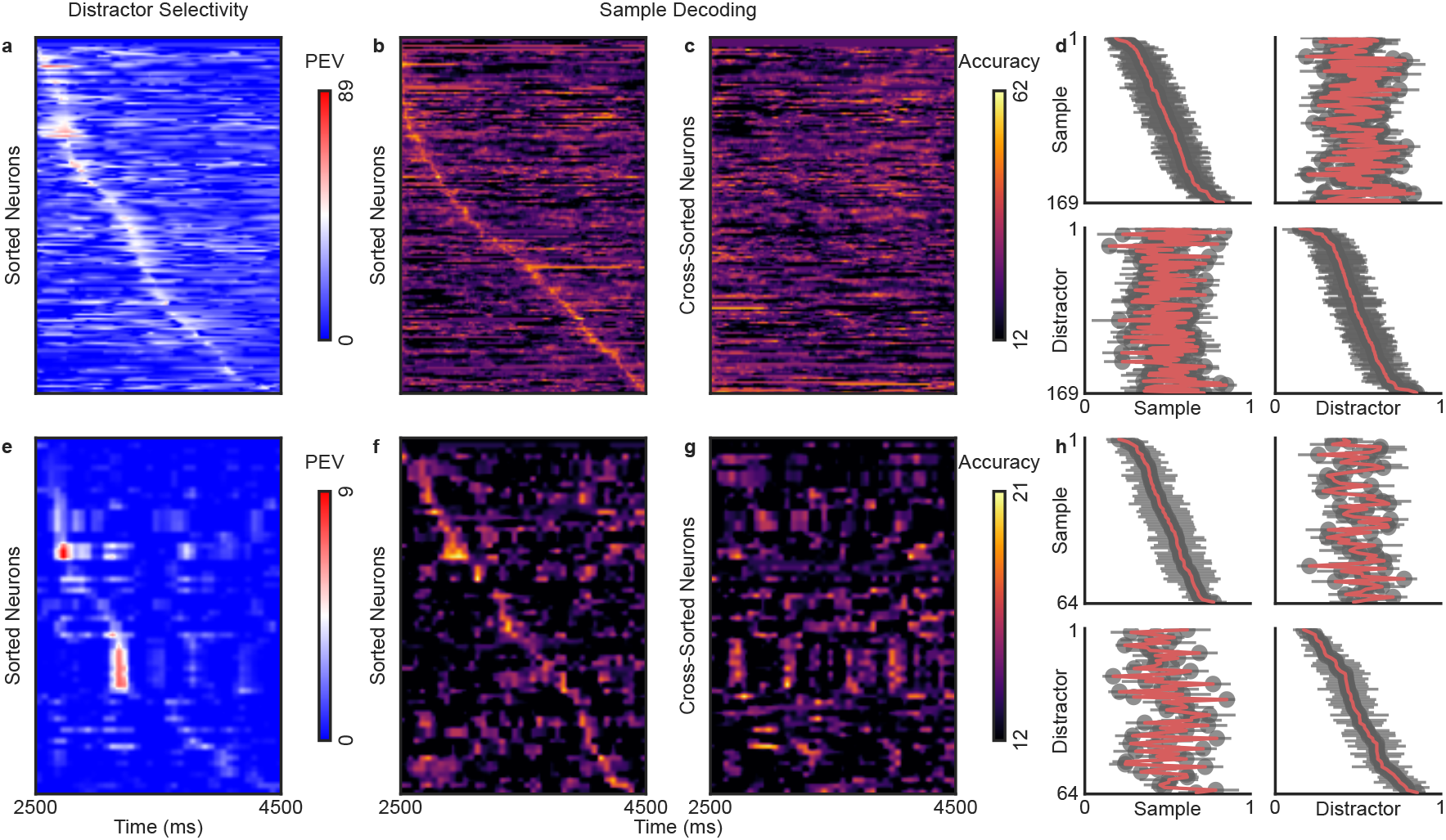
Working memory and distractor signals propagate along distinct pathways during distraction: Comparison between Forward-Biased RNN (a-d) and non-human primate (NHP) prefrontal cortex recordings (e-h). **(a, e)** Time-resolved distractor selectivity for each neuron (partial 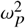; PEV from a two-way ANOVA), with neurons sorted by the latency of maximal distractor selectivity. **(b, f)** Time-resolved sample-decoding accuracy from a classifier trained on population firing rates; neurons are sorted by the latency of peak sample readout. **(c, g)** Cross-sorted sample-decoding maps in which neurons are ordered by their distractor-selectivity peak times (same orders as in panels a and e, respectively). In panels a-c and e-g, analyses show the interval from distractor onset (*t* = 2500 ms) to test onset (*t* = 4500 ms) in trials with the longest delay (*t*_*delay*_ = 4000 ms). In panels b, c, f, and g, the colorbar is adjusted to indicate the chance level in decoding accuracy. **(d, h)** Rank-stability across five delay durations. For each delay and measure (sample or distractor information), each neuron’s peak-time rank of the corresponding measure was normalized to [0,1] (x-axis) and compared to the mean rank across delays (y-axis). Within-factor comparisons (sample–sample, top–left; distractor–distractor, bottom–right) show strong stability, whereas cross-factor comparisons (sample–distractor and distractor–sample) lack structure, indicating distinct propagation orders. Error bars denote the s.e.m. of each unit’s normalized rank across delays.

To directly compare the temporal organization of sample-related and distractor-related activity, we cross-sorted the sample-decoding heatmap by each neuron’s distractor-selectivity peak latency. If the same neuronal sequence (i.e., the same order of neurons over time) governed both signals, the re-ordered sample-decoding heatmap would still exhibit a coherent cascade. Instead, this temporal structure in sample decoding broke down after cross-sorting in the Forward-Biased RNN (Fig. 4c). The same misalignment appeared in the NHP data (Fig. 4g). This verifies the model’s prediction that sample and distractor information were routed along overlapping but differently ordered sequences.

We next asked whether these sequences were consistent across varying delay lengths. We treated each of the five delay durations as a separate fold and computed the ranked peak time for each neuron for each factor. For each delay, we ranked neurons by the time of their peak response for each signal (sample decoding and distractor selectivity). We then normalized these ranks within each delay to the interval [0, 1]. For each neuron, we compared its normalized ranks with its mean rank across delays (Fig. 4d, h). This approach assesses whether a neuron’s relative position in the sequence is consistent across delays [63]. In the model, comparisons of ranks for sample decoding across delays revealed strong rank stability: neurons that carried sample information early in the sequence tended to do so across all delays, and likewise for late-peaking neurons (Fig. 4d, top-left). Distractor-selectivity of the neurons showed a similarly stable ordering across delays (Fig. 4d, bottom-right). The NHP recordings showed the same pattern of within-factor rank stability across delays for both sample and distractor signals (Fig. 4h, top left and bottom right).

Finally, we tested whether the propagation order of neurons carrying sample information was related to the propagation order of distractor responses across delays. In the Forward-Biased RNN, cross-factor comparisons (sample ranks versus distractor ranks) across delays showed no coherent structure (Fig. 4d, top-right and bottom-left), indicating no systematic relationship between the ordering of neurons in the sample sequence and in the distractor sequence. The NHP data displayed the same absence of coherent cross-factor structure (Fig. 4h, top-right and bottom-left).

Together, these results showed that both Forward-Biased RNN and NHP PFC recordings preserved the sample during distraction via a dynamic yet stable sequence of neurons. Meanwhile, the distractor was processed along a separate, equally stable sequence. Both pathways were consistent across delay lengths. This temporal and circuit-level segregation lets the system admit new input without overwriting stored memory.

## Discussion

Despite the prevalence of traveling waves in cortical circuits [7, 12], standard recurrent neural network (RNN) models do not exhibit such spatiotemporal patterns. Even when RNN units are arranged on a 2D sheet with spatially local, distance-dependent connectivity, these networks fail to support traveling waves. Here, we showed that introducing a forward bias into the locally connected recurrent circuitry produces and sustains traveling waves. Activity is relayed through one-way, feed-forward motifs rather than reciprocal loops. When the wave-enabled model was tested on a working memory task, information about the memory and a distractor intended to disrupt the memory traveled along segregated pathways. This was confirmed in neurophysiological data from NHP performing the same task.

This study addresses a fundamental question: can spatiotemporal traveling wave patterns emerge from fully trainable recurrent circuitry, without hand-crafted connectivity or synthetic neural dynamics? We employed a data-driven approach to induce traveling wave patterns in locally connected models. The circuit motifs supporting traveling waves emerged naturally by aligning RNNs to the neural manifold estimated from the neurophysiological data. Topological analysis revealed the network’s directed one-way connections were critical for generating waves, as they formed a latent feedforward chain of synaptic connectivity. Building on this insight, we incorporated these components *de novo* into the RNN architecture, alongside the spatial locality constraint. We integrated these elements by reparameterizing the recurrent weight matrix: A feedforward structure was introduced in a latent domain. It was then transformed to physical space via a similarity transform, where spatial locality was applied.

Computational models of traveling waves have garnered increasing interest across neuroscience and machine learning [16, 17, 24, 27, 64, 23, 46, 65, 66]. However, existing approaches predominantly rely on imposed dynamics or tailored connectivity. Wave-like motion is explicitly prescribed by equations, or by hand-designed connectivity templates applied uniformly across neuronal circuits. This reliance introduces a key limitation. Hard-coding dynamics or connectivity patterns in models can yield insight into how waves affect the models. However, they yield a limited understanding of how the waves emerge in the brain.

In contrast, we developed a theoretical framework where traveling waves emerge naturally through learning. The model includes soft, biologically motivated inductive biases but does not dictate dynamics or connectivity. We incorporated local connectivity as an anatomical prior [29, 30, 31], a constraint that has been shown to enhance biological plausibility [34, 67]. Then, we biased the recurrent weights toward a feed-forward structure in the latent space [57, 58], without hard-wiring connectivity patterns. Such directionally organized feed-forward motifs have been reported in electron-microscopy reconstructions of cortical microcircuits [68]. Consistent with this, Stroud et al. show that recurrent circuits with effectively feedforward dynamics can reproduce the temporal evolution of working memory codes observed in primate lPFC [69]. Despite these biases, synaptic weights in our model remain fully plastic. As a result, the network can deviate from the latent feedforward structure when task demands favor alternative configurations [58]. In this sense, the architecture is *wave-enabled* rather than *wave-imposed*.

While traveling waves have been implicated in cognitive operations, their computational function remains elusive [14, 46]. We found segregated paths for the propagation of working memories and a distractor. This segregation can limit interference between the two representations across space and time in the neural network, thereby protecting working memory. Oscillations may dynamically modulate neuronal selectivity (c.f. [70]). In doing so, they could help reallocate neural resources to encode evolving task demands. The observed dynamics may also reflect the spatiotemporal control mechanisms described by Spatial Computing Theory [71]. In this paradigm, oscillatory interactions route item-specific working memory representations across network space.

Neural circuits in the brain can leverage traveling waves for computation [1, 18, 28, 72]. However, this mechanism remains underused in modern artificial intelligence systems [73, 23, 24, 27]. Our model provides a computational framework for studying spatiotemporal computation in neural circuits. This framework offers a path to understand how coordinated wave activity may underlie cognitive function. Beyond cognition, wave-based analog computation can process information faster and more efficiently than digital code [73]. This paradigm could help overcome the limits of processor miniaturization and thermal dissipation. Moreover, such architectures could enable next-generation neuromorphic computing with cortical-like dynamics. Together, spatiotemporal neural computation models chart a promising path toward brain-inspired artificial intelligence.

## Methods

### Task description

As described in the Results, both the non-human primate recordings and the network simulations were performed on a delayed match-to-sample (DMS) task [10]. Each trial began with central fixation for 500 ms, followed by a 500 ms presentation of a *sample* stimulus drawn from a set of eight objects. The sample period was followed by a delay interval of duration *t*_delay_ ∈ {1000, 1410, 2000, 2830, 4000} ms. On half of the trials, a *distractor* stimulus, randomly selected from two objects that never appeared as sample items, was presented for 250 ms halfway through the delay. After the delay, two test stimuli were shown for 500 ms: the original sample (match) and a non-match randomly chosen from the remaining seven objects. Test items appeared at fixed coordinates (−0.5, 0.5) and (0.5, −0.5) within a [−1, 1] × [−1, 1] domain, with the match location randomly assigned on each trial. In the NHP experiments, the subject indicated its choice with a saccade to one of the two test locations. In the network simulations, the choice was read out as a two-dimensional response *y* corresponding to the selected item (see Network model below).

### Network model

In Fig. 1, we modeled population activity with a leaky continuous-time rate RNN whose hidden state ***z***_*t*_ ∈ ℝ^*N*^ evolves as

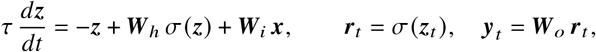

where *σ*(·) is a pointwise nonlinearity, ***W***_*i*_ ∈ ℝ^*N*×*M*^ and ***W***_*h*_ ∈ ℝ^*N*×*N*^ are the input and recurrent weight matrices, and ***W***_*o*_ ∈ ℝ^2×*N*^ is a linear readout that returns the two-dimensional choice coordinates ***y***_*t*_. In our implementation, we set *σ* = ReLU, and used a single recurrent layer with state dimension *N* = 169. The input dimensionality was *M* = 30, where inputs ***x***_*t*_ encoded both task events and stimulus identity, with item identities one-hot encoded separately for each location. The recurrent matrix ***W***_*h*_ was initialized orthogonally in the Manifold Aligned and Locally Connected models. In all models, ***W***_*i*_ and ***W***_*o*_ were initialized using the Xavier method [74].

### Model training

We trained the network model by minimizing a composite objective,

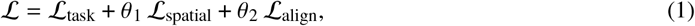

where **ℒ**_task_ quantifies task performance, **ℒ**_spatial_ encourages local connections, and **ℒ**_align_ promotes alignment with the target manifold. The coefficients *θ*_1_ and *θ*_2_ control the relative strength of each regularizing term.

Task performance was quantified by mean squared error (MSE) over output trajectories ***y***. The task loss across all trials was

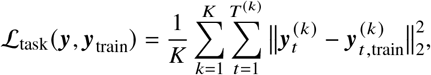

where *K* is the number of training sequences, *T* ^(*k*)^ is the length of trial *k*, and 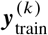 is the target output trajectory.

We assigned each neuron fixed coordinates on a square lattice. To encourage local connectivity, we regularized the magnitude of recurrent weights in proportion to the pairwise distances between neuronal positions,

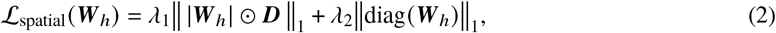

where *λ*_1_ and *λ*_2_ are regularization coefficients, ***D*** is the pairwise Euclidean distance matrix, and ⊙ denotes the Hadamard (element-wise) product. Because *D*_*ii*_ = 0, the distance-weighted term leaves self-connections unpenalized and could bias learning toward large autapses; the second term explicitly penalizes self-connections to prevent this.

To align latent population structure between model and PFC recordings, we constructed orthonormal bases ***Q***_RNN_ and ***Q***_NHP_ from the network’s hidden state and trial-averaged neural data, respectively, and computed the singular values *s*_*j*_ of 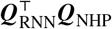. The alignment term maximized the sum of squared canonical correlations across the top four axes by minimizing

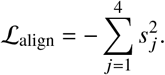

For training the Locally Connected baseline, the alignment term was not applied (*θ*_2_ = 0).

All models were implemented in PyTorch [75]. Network parameters were optimized with Adam [76] (learning rate 5 × 10^−4^ During training, only the input (***W***_*i*_) and recurrent (***W***_*h*_) weight matrices were updated. The linear readout (***W***_*o*_), which maps the recurrent state to the output, was held fixed at its Xavier initialization throughout optimization to ensure that learning was driven by the recurrent dynamics rather than the output mapping. After training, all model architectures achieved 100% correct performance on the DMS task.

### Forward-biased parameterization

In Fig. 3, we replaced the dense recurrent matrix with a structured similarity transform [58] that embeds a latent feed-forward scaffold,

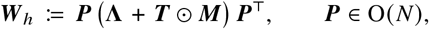

where *N* is the number of units. Here, ***T*** is a learnable strictly lower-triangular matrix that introduces forward-biased couplings, and ***M*** is a Gaussian mask that assigns greater weight to the immediate sub-diagonals, which dominate feed-forward interactions [54, 77, 57]. The matrix ***P*** is a learnable orthogonal change of basis (constrained via an exponential-map parametrization [78]), decoupling the latent feed-forward structure from the physical embedding used by the spatial prior.

Because a triangular matrix has eigenvalues equal to its diagonals (zero here) and orthogonal similarity preserves this spectrum, ***PTP***^⊤^ would place all eigenvalues at zero. To set the spectrum independently of ***T***, we introduce a separate learnable block-diagonal **Λ** [58] with 2 × 2 rotation-scaling blocks,

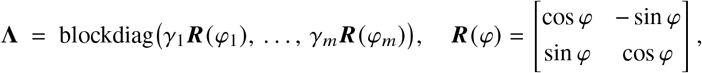

where *m* = ⌊*N*/2⌋. When *N* is odd, we append a 1 × 1 block as well. The gains *γ*_*i*_ and angles *φ*_*i*_ are learnable.

As in the baseline configuration, the Forward-Biased model was trained by minimizing Eq. 1. For this model, the alignment term was omitted (*θ*_2_ = 0).

### Wave dynamics

To visualize traveling waves in Figs. 1f and 3e, neurons were arranged on a square lattice, as assigned during training via the distance matrix ***D*** (Eq. 2). Hidden states were pooled with a Gaussian kernel (applied on a square window; width *h* = 5) to emulate the effect of the local field potential.

To quantify traveling wave dynamics, we estimated optical flow between consecutive time steps using the Horn–Schunck algorithm [35]. This approach reconstructs a velocity field that captures the underlying spatiotemporal wave patterns. For each trial sequence ***I***, we computed the instantaneous velocity field (*u, v*) at each frame ***I*** (*t*) by minimizing

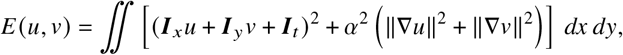

where ***I***_*x*_, ***I***_*y*_, and ***I***_*t*_ are spatial and temporal derivatives, and *α* is a smoothness coefficient (set to 15). The resulting linear system was solved iteratively with the Jacobi method, using a convergence threshold of *ϵ* = 10^−4^.

### Topological analysis

In Fig. 2, we computed graph-theoretic statistics on the network’s recurrent weight matrix ***W***_*h*_. For measures that require binary connectivity, we used proportional thresholding to obtain an adjacency matrix ***A***, retaining the 10% of edges with the largest absolute weights [34]. Analyses that required enumeration algorithms were performed using the Networkx package [79].

All plots show the mean outcome across 20 models, each initialized with a different random seed. The shaded regions in all statistics in Fig. 2, as well as in the alignment (Fig. 1e) and peak latency plots (Fig. 3a), indicate the 98% confidence interval, calculated via bootstrapping across models with 500 bootstrap samples.

#### Distance–weight correlation

In Figs. 2a and 3c, we reported the Pearson correlation *r* between the pairwise distance matrix ***D*** and the absolute recurrent weights |***W***_*h*_|,

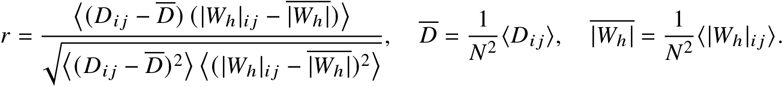

Here, ⟨·⟩ denotes the sum over all ordered neuron pairs (*i, j*).

#### Maximal cliques

A clique is a subset of vertices in which every pair of distinct vertices is connected by an edge. In other words, it is a fully connected subgraph. A clique is maximal if it cannot be extended by including any adjacent vertex, i.e., it is not contained within a larger clique. In Fig. 3b, we summarized how many maximal cliques of each size occur in the undirected graph ***Ã***. Let ℳ_*k*_ = {*C*: |*C*| = *k*} be the set of maximal cliques of size *k*. Figure 3b reports the histogram of counts 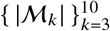. We used the Bron–Kerbosch algorithm [80] to find all maximal cliques.

#### Reciprocity

Figure 2c shows the reciprocity of the recurrent connection over the course of training. Reciprocity *p* is defined as the share of directed edges that have a counterpart in the opposite direction,

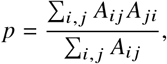

where *A*_*ij*_ = 1 indicates a directed edge from node *i* to node *j*.

#### Feed-forward loop ratio

In Fig. 2g, we used the Davis–Leinhardt directed triad census [81] to measure the fraction of closed triangles (where every node pair is connected by at least one directed edge) that are feed-forward loops (FFLs). If *n*_*τ*_ is the count of triads of type *τ*, the FFL ratio is the proportion of closed triads that are FFLs,

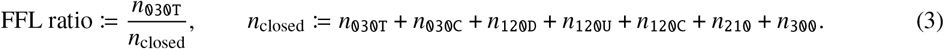

In this coding, `030T` is the feed-forward loop (*i* → *j, j* → *k, k* →*i*).

#### Directed 3-cycle ratio

Figure 2e reported the ratio of triads with directed 3-node cycles *n*_3-cyc_ in the graph. A directed 3-node cycle consists of three distinct nodes (*i, j, k*) such that there are directed edges from *i* to *j, j* to *k*, and *k* to *i*, forming a closed directed loop. Given the adjacency matrix ***A***, we compute

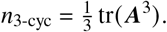

Each directed 3-node cycle contributes three closed walks, so we divided by 3 to avoid overcounting walks starting at different nodes. The cycle ratio is then computed as

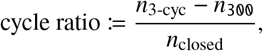

where *n*_closed_ is defined in Eq. 3. If all three edges are reciprocal (*τ*: `300`), the two possible orientations are counted separately. By subtracting *n*_300_, we ensured these are counted only once in both the numerator and denominator.

### Sample decoding

To quantify how much information about the sample each neuron carries over time, we used a time-resolved multiclass decoding analysis on single-unit activity. In each time bin, a multinomial logistic regression predicted the sample identity from the spike rate activity of a single neuron. To obtain reliable estimates, we used stratified *K*-fold cross-validation, with *K*=2 for the network model (two trials per sample) and *K*=5 for the neural recordings. Stratification preserved the label composition in every fold (by ‘sample × distractor’ for the recordings, and by ‘sample’ for the model). Decoding performance is reported as balanced accuracy (i.e., the mean recall across sample classes), averaged across folds. This produces a unit-by-time sample-decoding map for each session or model. Chance performance for eight classes is 1/8 (12.5%); the minimum of the colorbars in Fig. 4b-c,f-g is set to this level.

### Distractor selectivity

Selectivity was quantified with a *two-way* ANOVA applied to single-unit activity in each time bin, with factors *sample* and *distractor*. Type-II sums of squares were used so that the contribution of each factor is evaluated after accounting for the other. For *distractor* factor, we report the partial omega-squared (a bias-corrected proportion of explained variance, PEV [62]) as

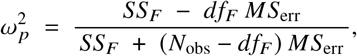

where *SS*_*F*_ and *df*_*F*_ are the sum of squares and degrees of freedom for factor, *MS*_err_ is the mean squared error, and *N*_obs_ is the number of trials.

## Acknowledgments

N.S.T. thanks Robert Rosenbaum and Tamal Batabyal for their helpful discussions. The authors thank Jordan DeFarias for assistance with data collection. This work was supported by Army Research Office grant W911NF2410228, Office of Naval Research (ONR) MURI grant N00014-23-1-2768, Air Force Office of Scientific Research (AFOSR) grant FA9550-21-1-0223, Freedom Together Foundation, the Picower Institute for Learning and Memory, and by the Swartz Foundation Fellowship for Theory in Neuroscience to N.S.T.

## Notes

### Competing Interest Statement

The authors have declared no competing interest.

