## Supplementary Materials for "Emergent Traveling Waves in Neural Circuits"

### Supplementary Material

#### Post-distractor activation decay

Figure S1 compares firing rate patterns across RNN architectures from distractor onset through the end of the delay period. This comparison revealed two distinct dynamical regimes. In Standard and Locally Connected RNNs, the population response to the distractor showed a rapid rise in activity, followed by monotonic decay to baseline. This pattern is consistent with diffusive relaxation. In the Locally Connected RNN, this relaxation manifested as a collapse of domain-averaged wave strength, as quantified by optical-flow analysis (Fig. 1). In contrast, Manifold-Aligned and Forward-Biased RNNs exhibited a sequentially propagating cascade throughout the post-distractor delay. This cascade is mirrored by persistent traveling waves (Fig. 3). These activity patterns distinguish the latter models from their standard and Locally Connected counterparts.

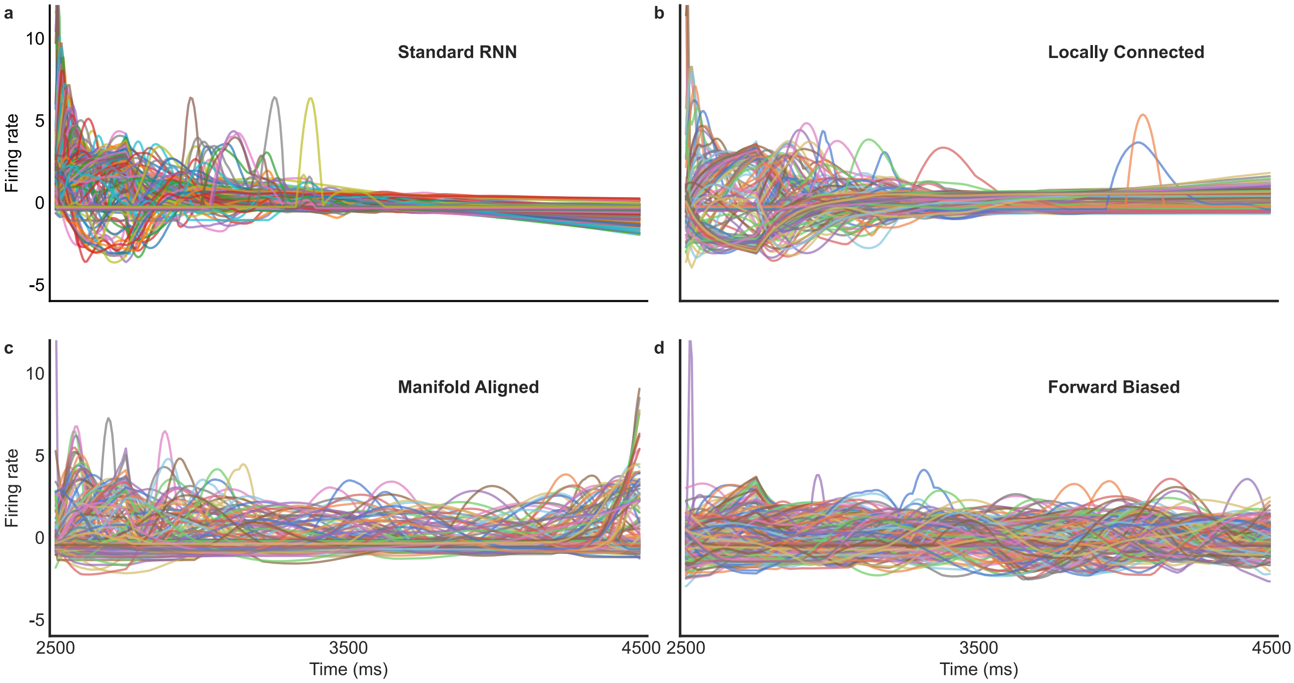


Figure S1: **The firing rates in the Locally Connected model decays to a nearly persistent level during the post-distraction delay:** Comparing trial-averaged firing rate patterns across models during distraction in the delayed match-to-sample (DMS) task, for the longest delay length $t_{delay}$ = 4000. **a,** standard RNN **b,** Locally Connected **c,** Manifold-Aligned **d,** Forward-Biased models. Colored traces show single-unit firing rates.

#### Post-distractor firing rate histograms

To complement the spatial analyses of wave propagation presented in the results, we illustrate peri-event time histograms (PETHs) of spike rate activity across the neuronal population during distraction (Fig. S2). In this representation, a diagonal ridge in the peak-ordered PETH emerges as the canonical signature of a traveling wave through the population [64, 28]. For each model and recording, trial-averaged spike rate activity was computed separately for each delay condition and subsequently z-scored across time for each neuron. Each panel depicts the temporal evolution of population activity from distractor onset through the end of the delay period. In all cases, the distractor was displayed during the initial 250 ms of the interval. Analyses of both NHP recordings and wave-enabled RNNs revealed a pronounced post-distractor diagonal band of activity. Notably, this band systematically propagated from early- to late-peaking units and recurred reliably across variable delay durations.

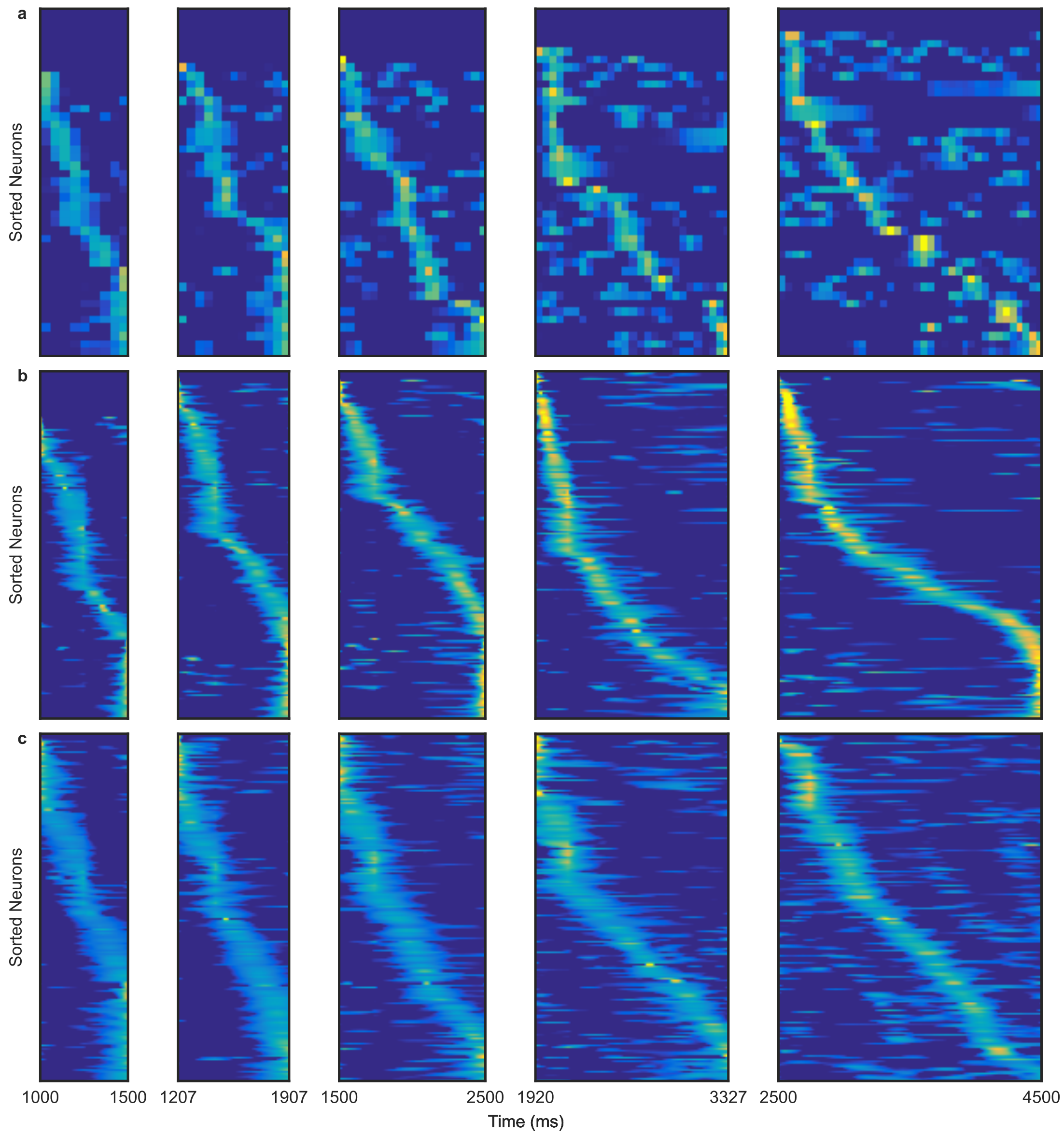


Figure S2: **Post-distractor spike rate activity reveals traveling wave sequences that scale with delay in cortex and wave-enabled RNNs:** Peri-event time histograms during distraction for **a,** non-human primate recordings; **b,** Manifold-Aligned model; and **c,** Forward-Biased model. Heat maps display trial-averaged spike rate activity over time (x-axis). Neurons are ordered by the latency of their peak activity. Each panel begins at the mid-delay distractor onset and ends at the delay offset. Columns represent increasing delay durations from left to right.

#### Distance-dependence in Locally Connected model

To assess the distance dependence of connectivity in the base Locally Connected model following training, we quantified the relationship between the magnitude of recurrent weights $|W_h|_{ij}$ and the physical separation of units $D_{ij}$ on the 2-D sheet (see Methods, Eq. 2). In the Locally Connected network, the mean absolute recurrent weight showed a pronounced decrease as pairwise distance increased (Fig. S3). Stronger synaptic connections were found among spatially proximal units. In contrast, long-range connections tended to be weaker on average.

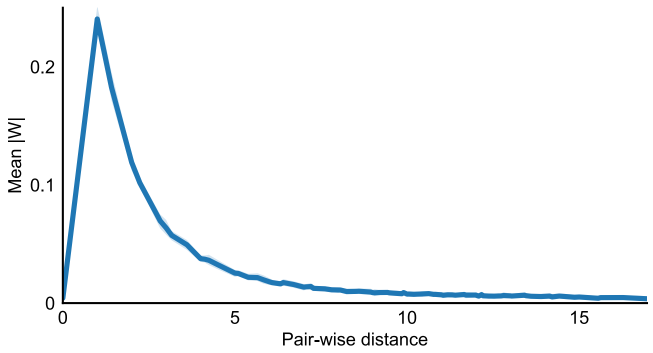


Figure S3: **Distance-dependent regularization promotes local connectivity:** Distance dependence of recurrent connections in the trained Locally Connected network. Mean absolute value of recurrent weights $|\boldsymbol{W}_h|$ (y-axis) is shown as a function of pair-wise distance between units on the cortical sheet $\boldsymbol{D}$ (x-axis). The shaded region illustrates the 98% confidence interval.

#### Standing waves in the Locally Connected model

In Figs. 1 and 3, we showed that the Manifold-Aligned and Forward-Biased models exhibit traveling waves during the working memory delay. By contrast, the Locally Connected model (constrained only to have stronger synaptic connections between spatially proximal neurons) did not produce traveling waves. Instead, this architecture gave rise to standing wave patterns. Standing waves are spatially organized activity patterns whose amplitudes vary over time while their spatial profiles remain stationary (i.e., they do not propagate across the network). Figure S4 demonstrates the observed patterns in the Locally Connected model for a representative trial with the shortest delay ($t_{\text{delay}}$ = 1000 ms). We observed two distinct standing waves: the first was evoked by the sample stimulus (presented at $t = 0 - 500$ ms), and the second was triggered by the distractor stimulus (presented at $t = 1000 - 1250$ ms). Local connectivity strengthened interactions among neighboring neurons, producing smooth spatial changes of activity. However, local connectivity alone was not sufficient to support wave propagation, so the resulting patterns remain spatially localized.

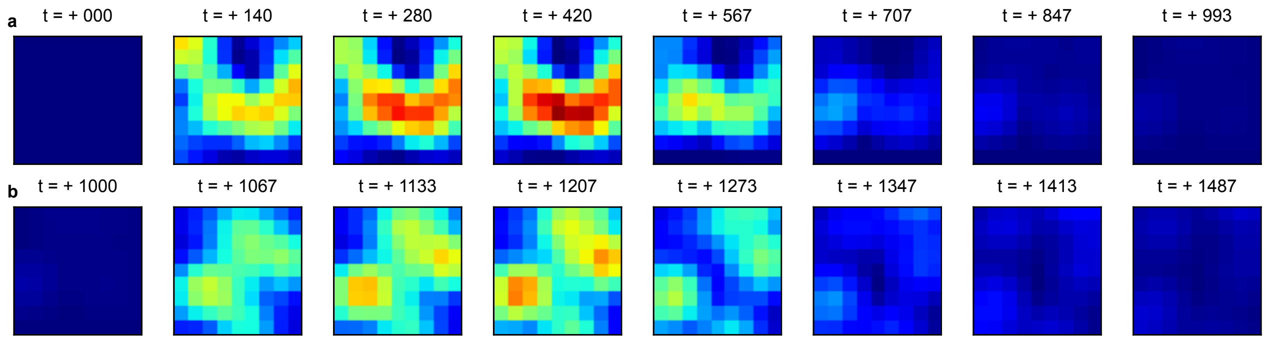


Figure S4: **The Locally Connected model generates stimulus-evoked standing-wave patterns in neural population activity:** Heatmaps illustrate wave patterns across the model's spatially arranged neurons from a representative trial with $t_{\text{delay}}$ = 1000 ms. **a,** Wave patterns during sample stimulation ($t = 0 - 500$ ms) and the subsequent delay period ($t = 500 - 1000$ ms). **b,** Wave patterns during distractor stimulation ($t = 1000 - 1250$ ms) and the following delay ($t = 1250 - 1500$ ms).

#### Traveling waves observed in empirical data during memory delay

To visualize traveling waves in the empirical data, we analyzed local field potentials (LFPs) recorded from the PFC of an NHP. The LFP data were collected using the same multielectrode arrays and the same delayed match-to-sample sessions as those used to estimate the empirical neural manifold (Fig. 1c, e). Figure S5 displays a sequence of spatial snapshots of LFP activity from a representative trial with the longest delay ($t_{delay}$ = 4000 ms). The memory delay began at $t$ = 500 ms and ended at $t$ = 4500 ms, with a brief visual distractor presented from $t = 2500$ to $t = 2750$ ms. The spatial pattern of LFP activity formed a coherent wavefront that shifted across the array over time, consistent with propagating traveling waves. These results demonstrate that robust traveling waves were present during working memory maintenance in the absence of external sensory input, mirroring the model dynamics described in the main text.

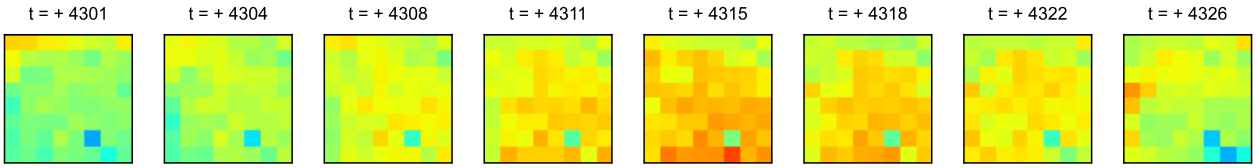


Figure S5: **Traveling waves are present in neural activity recorded from NHP PFC:** Sequence of spatial maps of local field potential (LFP) activity recorded from PFC on a multielectrode array during a representative trial with the longest delay ($t_{delay}$ = 4000 ms).

#### Propagation of information in the Manifold-Aligned model

In the main text, we showed that working memory and distractor information propagated along distinct pathways during the distraction and subsequent delay periods. We showed this in the Forward-Biased model and in electrophysiological recordings. Here, we verified that this sequential information propagation is also present in the Manifold-Aligned networks. For this model, we computed time-resolved distractor selectivity for each neuron (partial $\omega_p^2$; PEV from a two-way ANOVA) from spike rate neural activity. We then computed time-resolved sample-decoding accuracy using a classifier trained on population firing rates (see Methods). Next, we sorted neurons separately by the latency of their peak value for each delay period and each measure (sample or distractor information). We then compared these ranks across delays and measures to determine whether they were consistent across delay periods and how they related across the two factors (sample vs. distractor; Fig. S6). The Manifold-Aligned network exhibited a sequential organization: units that reached peak sample decoding earlier in the sample sequence did so consistently across all delay lengths, and units that peaked later in the sequence maintained their relative temporal positions across delays. A similar pattern was observed for distractor-related information. Notably, cross-factor analyses revealed that the ordering of units for sample and distractor information was not preserved between factors. This suggests that, within the Manifold-Aligned model, sample and distractor information propagated via distinct neural pathways.

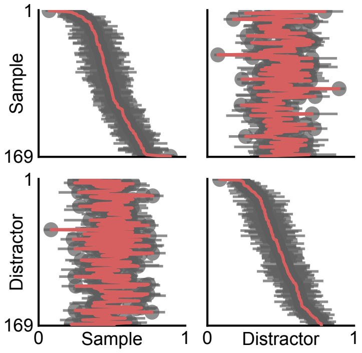


Figure S6: **Working memory and distractor information propagate along distinct pathways in the Manifold-Aligned network:** Rank stability across five delay durations for the Manifold-Aligned RNN. For each delay and measure (sample or distractor information), each neuron's peak-time rank was normalized to [0,1] (x-axis) and compared to its mean rank across delays (y-axis). Within-factor comparisons (sample–sample, top left; distractor–distractor, bottom right) show strong stability, whereas cross-factor comparisons (sample–distractor and distractor–sample) lack structure, indicating distinct propagation orders. Error bars denote the s.e.m. of each unit's normalized rank across delays.
